# Human B cell lineages engaged by germinal centers following influenza vaccination are measurably evolving

**DOI:** 10.1101/2021.01.06.425648

**Authors:** Kenneth B. Hoehn, Jackson S. Turner, Frederick I. Miller, Ruoyi Jiang, Oliver G. Pybus, Ali H. Ellebedy, Steven H. Kleinstein

## Abstract

Poor efficacy of seasonal influenza virus vaccines is often attributed to pre-existing immunity interfering with the persistence and maturation of vaccine-induced B cell responses.^1^ Consistent with this notion, no significant increase in somatic hypermutation (SHM) among circulating influenza-binding lineages was detected following seasonal vaccination in humans.^2^ A more recent study showed that at least a subset of vaccine-induced B cell lineages are recruited into germinal centers (GCs) following vaccination, suggesting that affinity maturation of these lineages can occur.^3^ Crucially, however, it has not been demonstrated whether these GC-engaged lineages actually accumulate additional SHM. Here, we address this point using a phylogenetic test of measurable evolution. We first validate this test through simulations and demonstrate measurable B cell evolution in known examples of affinity maturation such as the response to HIV infection. We then show that lineages in the blood are rarely measurably evolving following influenza vaccination, but that GC-engaged lineages - likely derived from memory B cells - are frequently measurably evolving. These findings confirm that seasonal influenza virus vaccination can stimulate additional SHM among responding B cell lineages.

## Main text

Measurably evolving populations are evolutionary systems that undergo mutation and selection rapidly enough to be detected in longitudinally-sampled timepoints.^4^ While this concept is frequently applied to viruses such as HIV^5^ and SARS-CoV-2,^6^ B cells experience similarly rapid evolution during affinity maturation. B cell affinity maturation is critical for developing high affinity antibodies in response to infection and vaccination.^7^ During affinity maturation, somatic hypermutation (SHM) introduces mutations into the B cell receptor (BCR) loci orders of magnitude more rapidly than the background rate of somatic mutations.^8^ These modified BCRs are selected based on their binding affinity, and the process repeats cyclically within GCs.^7,9^ Infection or vaccination can also stimulate pre-existing memory B cells that rapidly differentiate into antibody secreting plasmablasts and rarely re-enter GCs to undergo additional affinity maturation.^1,10^ A lack of vaccine-specific affinity maturation is thought to underlie the poor efficacy of seasonal influenza virus vaccination.^1,11^ While recent work has shown that antigen-specific B cell lineages can be recruited into GCs following influenza vaccination,^3^ it is not known whether these lineages then accumulate additional SHM.

Whether seasonal influenza vaccination stimulates additional SHM can be answered by determining whether GC-engaged, influenza-binding B cell lineages are measurably evolving following vaccination. This is distinct from simply quantifying SHM frequency. While influenza vaccination stimulates memory B cell lineages with high SHM frequency,^12,13^ these lineages are only measurably evolving if they undergo additional SHM during the sampling interval surrounding vaccination. In this study, we show how a phylogenetic test of measurable evolution can be a powerful tool to detect ongoing SHM in longitudinally-sampled BCR sequence datasets.^14,15^ We validate this approach through simulations and a survey of measurable evolution in B cell repertoires across a wide range of infections and vaccinations. We document significant heterogeneity among conditions, with some like HIV infection and primary hepatitis B vaccination enriched for measurably evolving lineages in the blood. We further show that while most circulating lineages following influenza virus vaccination are not measurably evolving, a subset of memory B cell lineages re-enter GCs and rapidly gain additional SHM.

Testing for measurable evolution in B cells requires longitudinally sampled sequence data from the BCR variable region. After pre-processing, we identify clonal lineages – B cells that descend from a common V(D)J rearrangement – using clustering based on nucleotide sequence similarity, which we have previously shown detects clonal relationships with high confidence.^16,17^ The pattern of shared SHM among BCR sequences within a lineage is used to build a maximum parsimony tree, which represents a lineage’s history of SHM. Branch lengths within these trees represent SHM/site. The divergence of each tip is the sum of branch lengths leading back to the most recent common ancestor. In evolving lineages, sequences at later timepoints are expected to have higher divergence than those from earlier timepoints (**Fig. 1a**). To estimate the rate of evolution over time, we calculate the slope of the regression line between timepoint (weeks) and divergence (SHM/site) for each tip (**Fig. 1b, e**).^18^ Because tips are not independent, standard linear-regression p values are improper. We instead quantify significance using a modified phylogenetic date randomization test.^14,15^ This tests whether the Pearson correlation between divergence and time is significantly greater than that observed in the same tree with timepoints randomized among tips (**Fig. 1c, f**).^14,15^ To account for population structure and sequencing error, we permute timepoints among single-timepoint monophyletic clusters of tips rather than individual tips (**Supplemental Figs. 1, 2**).^14,15^ We refer to lineages with a date randomization test p < 0.05 as “measurably evolving.” To limit our analyses to lineages with adequate statistical power, we include only lineages with ≥15 total sequences sampled over at least three weeks, and have a minimum possible p value < 0.05 based on the number of distinct permutations. Because we use a p value cutoff of 0.05, we expect a false positive rate of approximately 5% if no measurable evolution is occurring. We therefore refer to datasets with >5% measurably evolving lineages as “enriched” for measurable evolution. This test is implemented within the Immcantation.org framework in the R package *dowser*.^19^

**Fig. 1:**
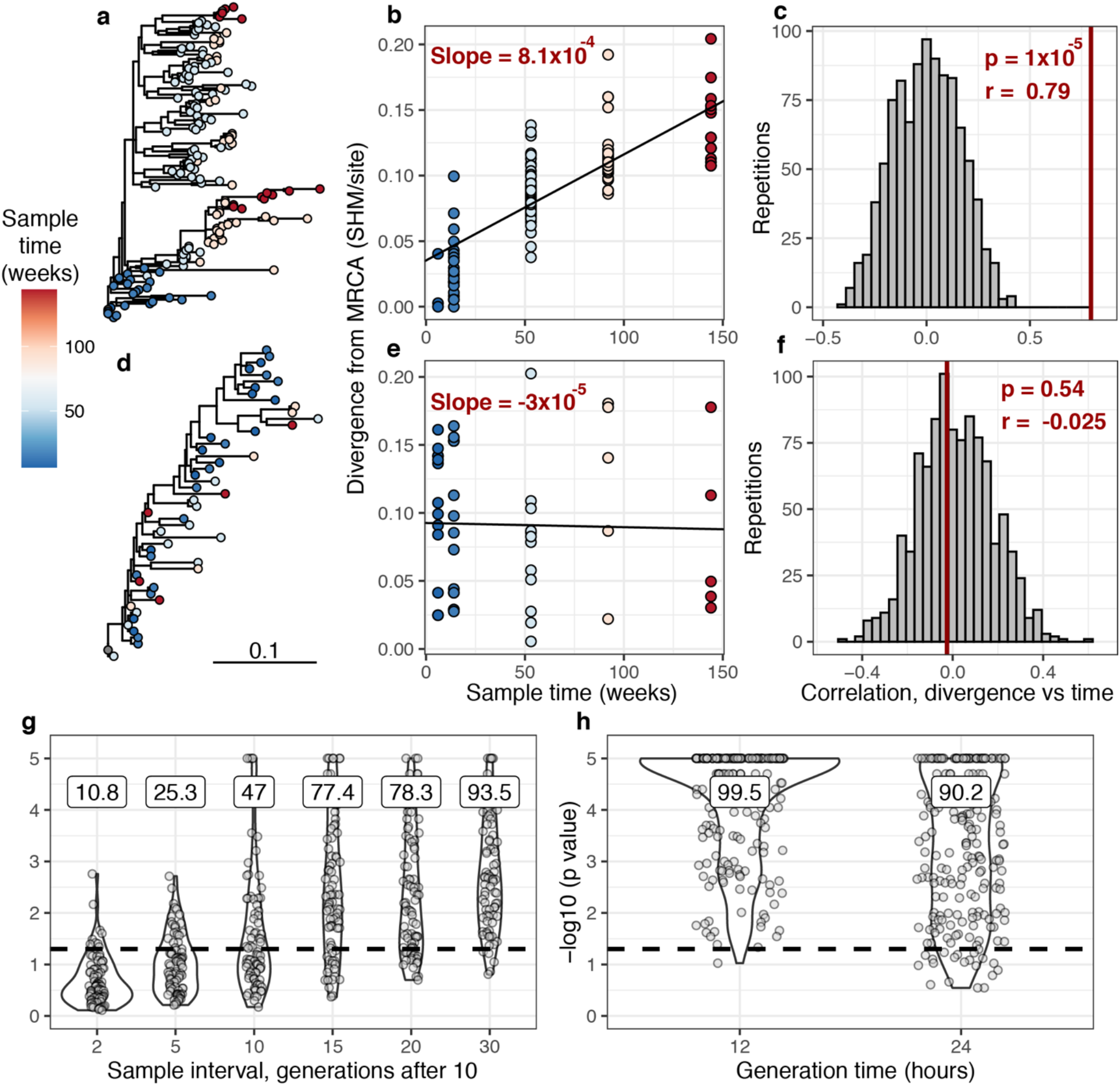
Detecting measurable evolution in B cell lineages. **a)** Example B cell lineage tree showing increasing divergence with sample time. Branch lengths show SHM/site according to scale bar in **d**. **b)** Rate of SHM accumulation over time estimated using a regression of divergence vs time in tree **a**. **c)** Significance of the relationship between divergence and time estimated using a date randomization test comparing the Pearson’s correlation (r) between divergence and time in tree **a**. **d-f)** Same plots as **a-c** but on a tree that is not measurably evolving. **g)** Simulation-based power analysis shows permutation test has high power over an interval of at least 10-30 GC cycles (generations). Lineages were sampled once at generation 10, and a second after the specified number of additional generations have elapsed. Percentage of lineages with p < 0.05 are listed above, rounded to three significant digits. The dotted line corresponds to p = 0.05. **h)** Simulation-based analysis reproducing the sampling of Laserson et al. (2014)^13^ shows the test has high power even at slow (24 hour) GC cycle times.

To determine the necessary sampling interval to detect ongoing SHM, we benchmarked the date randomization test using affinity maturation simulations performed with the package *bcr-phylo*.^20,21^ This simulates alternating GC cycles of B cell proliferation, SHM, and selection based on amino acid similarity to a target sequence. Within these simulations, each lineage was first sampled after 10 simulated GC cycles, and then sampled a second time after a variable number of additional cycles. Using this framework in which all lineages are evolving, the date randomization test detected measurable evolution in 47% of lineages after 10 additional GC cycles, and 77% after 15 additional cycles (**Fig. 1g**). Given a GC cycle time of 6 – 24 hours, 15 cycles corresponds to 4 – 15 days, within the timeframe of many longitudinal B cell repertoire studies.^2,13^ To quantify the false positive rate, we repeated these calculations on the same simulations but with randomized sample time associations. Here, the date randomization test found measurable evolution in < 2.5% of cases, indicating a low false positive rate (**Supplemental Fig. 3**). These analyses demonstrate that the date randomization test has sufficient sensitivity and specificity to detect ongoing SHM from longitudinally sampled BCR data.

To further validate our approach, we tested for measurable evolution in cases of known or suspected affinity maturation in humans. We hypothesized that primary immune responses would be enriched for measurably evolving lineages.^8^ To test this, we used publicly available data primarily from the Observed Antibody Space database^22^ to survey measurable evolution in BCR datasets from 99 human subjects in 21 studies spanning 10 conditions including HIV infection, Ebola virus infection, and healthy controls (**Table 1**). We observed considerable heterogeneity in measurable evolution among conditions. Confirming our hypothesis, we observed an enrichment of measurably evolving lineages (>5% of tested lineages) in primary immune responses including HIV infection, meningococcus vaccination, primary but not secondary hepatitis B vaccination, and early childhood development (**Table 1; Fig. 2a**).

**Table 1:** Summary of datasets. *Subjects* shows number of subjects with at least one powered lineage. *Mean range* shows mean total sampling interval across subjects. *Powered lineages* shows the number of lineages that: i) contained at least 15 sequences, ii) were sampled over at least 3 weeks, and iii) had a minimum possible p value < 0.05. The rightmost column shows the percentage of these lineages with p < 0.05, rounded to two significant digits. Studies with > 5% positive lineages are shown in bold. Turner et al. (2020) in this table and **Fig. 2** included only blood samples. Data from studies marked with an asterisk (*) were obtained from Observed Antibody Space.^22^

| Study | Condition | Subjects | Mean range (weeks) | Mean sample count | Multi-timepoint lineages | Powered lineages | % lineages $p < 0.05$ |
| --- | --- | --- | --- | --- | --- | --- | --- |
| Levin et al. (2015)*<sup>33</sup> | Allergy + SIT | 9 | 52 | 2.7 | 42 | 31 | **6.5** |
| Davis et al. (2019)*<sup>32</sup> | Ebola virus | 4 | 36 | 3.6 | 1549 | 869 | 4.4 |
| Wang et al. (2014)<sup>43,44</sup> | Healthy adults | 7 | 52 | 2 | 18 | 10 | 0 |
| Nielsen et al. (2019)<sup>28</sup> | Healthy children | 20 | 69 | 2.7 | 262 | 71 | **14** |
| Galson et al. (2015a)*<sup>30</sup> | Hep. B vaccine | 9 | 4 | 4.8 | 4923 | 3422 | 2.9 |
| Galson et al. (2016)*<sup>31</sup> |  | 9 | 23 | 6.9 | 4426 | 2529 | **7.2** |
| Doria-Rose et al. (2014)*<sup>25</sup> | HIV | 1 | 190 | 8 | 65 | 48 | **44** |
| Huang et al. (2016)*<sup>58</sup> |  | 1 | 120 | 12 | 388 | 221 | **5.9** |
| Johnson et al. (2018)*<sup>59</sup> |  | 1 | 170 | 5 | 561 | 330 | **23** |
| Landaïs et al. (2017)*<sup>60</sup> |  | 1 | 160 | 7 | 1084 | 743 | **48** |
| Liao et al. (2013)*<sup>24</sup> |  | 1 | 140 | 5 | 205 | 151 | **53** |
| Schanz et al. (2014)*<sup>61</sup> |  | 1 | 120 | 3 | 147 | 54 | **11** |
| Setliff et al. (2018)*<sup>62</sup> |  | 6 | 170 | 3 | 787 | 173 | **9.8** |
| Wu et al. (2015)*<sup>26,63</sup> |  | 1 | 730 | 7 | 393 | 305 | **26** |
| Ellebedy et al. (2016)*<sup>2</sup> | Influenza vaccine | 8 | 13 | 5 | 1966 | 1479 | **5.2** |
| Laserson et al. (2014)*<sup>13,17</sup> |  | 3 | 4 | 9 | 1182 | 639 | 4.9 |
| Turner et al. (2020)<sup>3</sup> |  | 1 | 8.6 | 5 | 168 | 104 | 2.9 |
| Galson et al. (2015b)*<sup>29</sup> | Meningococcus vaccine | 7 | 4 | 3 | 483 | 80 | **10** |
| Jiang et al. (2020a)<sup>64</sup> | Myasthenia gravis | 3 | 260 | 3.6 | 110 | 62 | 3.2 |
| Jiang et al. (2020b)<sup>65</sup> |  | 1 | 52 | 2 | 46 | 33 | 3 |
| Tsioris et al. (2015)<sup>66</sup> | West Nile virus | 6 | 5.2 | 2 | 151 | 65 | 1.5 |

**Fig. 2:**
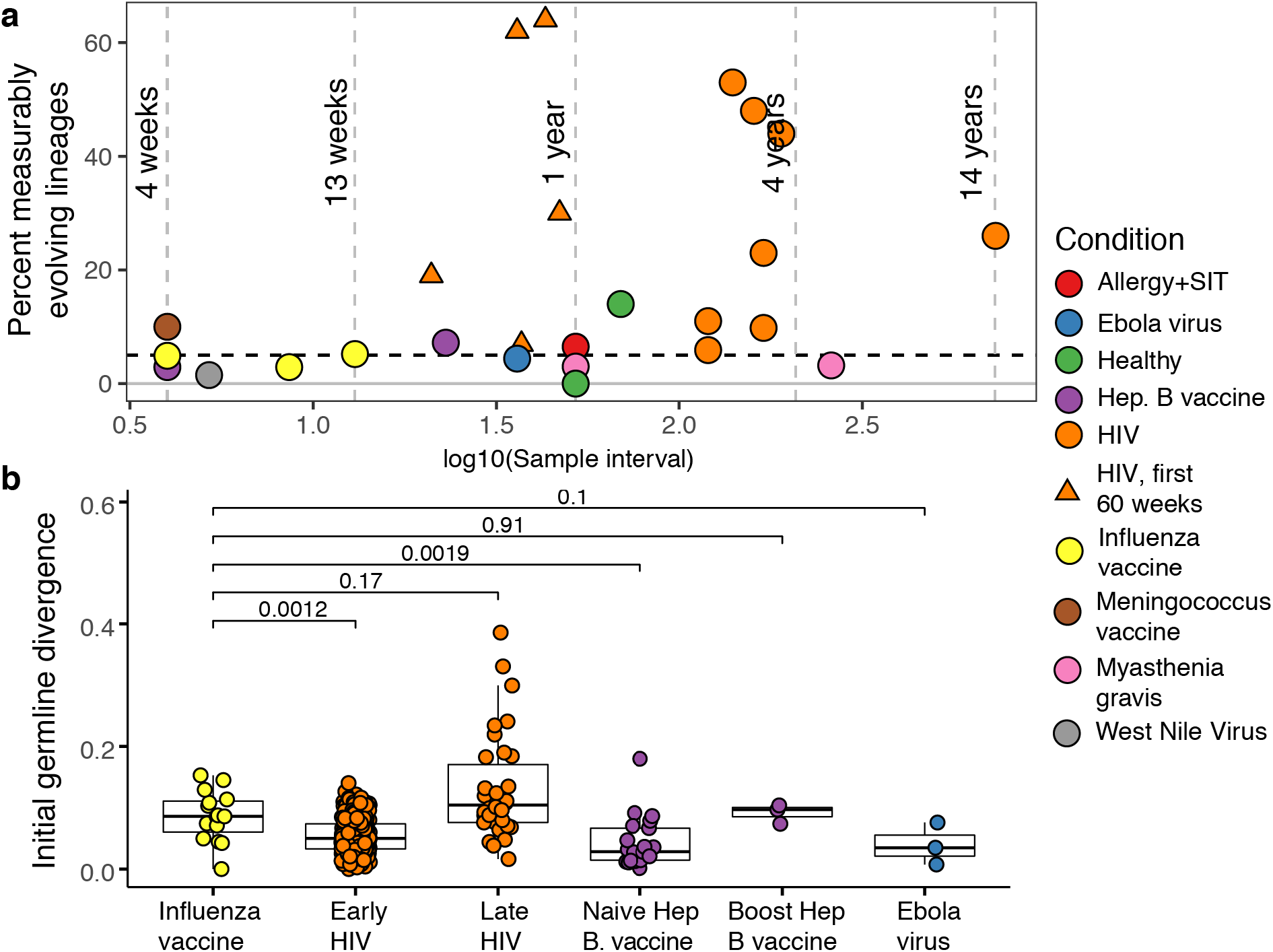
Measurable evolution in B cell lineages across time and conditions. **a)** Percentage of lineages that are measurably evolving within each study (**Table 1, Fig. 1c**). The dotted line indicates 5%, the percent expected under the null hypothesis that there is no measurable evolution occurring in a given dataset. Triangles indicate HIV datasets down-sampled to only include samples before 60 weeks. **b)** Mean divergence (sum of branch lengths) from germline to sequences from each adjusted measurably evolving lineage’s first timepoint. Each point is a measurably evolving lineage with a Benjamini-Hochberg adjusted p value < 0.1. Wilcoxon tests were used to compare divergence levels among datasets.

Chronic HIV infection stimulates ongoing affinity maturation as B cells evolve to contain viral escape mutants.^23,24^ Consistent with this arms race, HIV infection was more enriched for measurably evolving lineages than other conditions surveyed, with between 5.9% and 53% of lineages measurably evolving (**Fig. 2a**). Lineages from patients with broadly neutralizing anti-HIV lineages sampled over multiple years^24–27^ were particularly enriched (26% – 53% measurably evolving). Importantly, the HIV studies included were sampled over longer time periods than studies of other conditions (mean = 225 vs 45 weeks; **Table 1**). To determine whether these results were simply due to longer sampling intervals, we repeated our analysis of HIV patients using only samples within the first 60 weeks. These truncated datasets were still highly enriched for measurably evolving lineages (6.9% – 64%) compared to other datasets with similar sampling intervals (0% – 7.2%, **Fig. 2a**). This indicates that the observed high frequency of measurably evolving lineages is not simply due to long sampling intervals.

Other primary immune responses were also enriched for measurably evolving lineages (**Table 1**, **Fig. 2a**). B cell lineages from healthy children sampled during the first three years of life were enriched for measurable evolution (14%), possibly reflecting continual exposure to novel antigens.^28^ We also observed an enrichment of measurably evolving lineages following primary meningococcus vaccination (10%)^29^ and primary but not secondary hepatitis B vaccination (7.2% vs. 2.9%, respectively).^30,31^ Primary hepatitis B vaccinees were sampled over a longer time period than secondary vaccines, so this difference may also be due to different sampling intervals (**Supplemental Fig. 5**). Interestingly, Ebola virus infection showed a low (4.4%) percentage of measurably evolving lineages (**Table 1**) despite likely being a primary infection. However, 2/14 (14.3%) lineages containing Ebola virus specific monoclonal antibodies were enriched for measurable evolution, suggesting Ebola virus binding lineages were evolving during the study.^32^ Finally, allergen-specific immunotherapy, which stimulates tolerance of allergy-causing antigens through exposure, was also enriched for measurable evolution (6.5%).^33^ Overall, these results confirm that the date randomization test can detect ongoing SHM in datasets where it is expected to be occurring.

Seasonal influenza vaccination is believed to trigger a memory B cell response in adults. If memory B cells rarely re-enter GCs to undergo additional affinity maturation, we expect little measurable evolution in the blood following vaccination.^10^ To test this, we applied the date randomization test to three longitudinally-sampled adult influenza vaccine datasets. The first comprised 3 adults sampled 7 times between 1 hour and 28 days post-vaccination^13,17^; the second contained 8 adults sampled 5 times between 0 and 90 days post-vaccination^2^; the third used blood samples from a single individual sampled 5 times between 0 and 60 days post-vaccination.^3^ Across all subjects in each study, between only 2.9% and 5.2% of lineages were measurably evolving (**Table 1**). These values are approximately as expected under the null hypothesis of no measurable evolution, and histograms of p values from these datasets are roughly uniform, suggesting the measurably evolving lineages identified are mostly false positives from multiple testing (**Supplemental Fig. 6**). To verify the 4 – 13 week sampling range of these studies was sufficient to detect measurable evolution, we performed simulation analyses replicating the sampling strategy of the influenza dataset with the shortest sampling range (**Supplemental Fig. 4**).^13^ These simulations show this timescale was sufficiently long to detect ongoing affinity maturation with high sensitivity (>90%, **Fig. 1h**). Overall, these results indicate B cell lineages present in blood infrequently undergo additional evolution within 13 weeks following influenza vaccination, consistent with a primarily GC-independent memory B cell response and/or rarity of antigen-specific lineages in the peripheral blood.^12^

While measurably evolving lineages were not enriched in the blood following influenza vaccination, we checked if any could be identified after adjustment for multiple testing. We identified 15 lineages in influenza datasets, and 342 lineages in other conditions, with Benjamini-Hochberg^34^ adjusted date randomization p values < 0.1. We investigated if these “adjusted” measurably evolving lineages were derived from naive or pre-existing memory B cells. Because memory B cell lineages have already undergone affinity maturation, they will have higher initial SHM levels compared to naive B cell lineages. To test this, we compared germline sequence divergence in adjusted measurably evolving lineages from influenza vaccination to other conditions. Consistent with memory B cell re-activation, lineages from influenza vaccination had significantly higher initial divergence (median = 8.6%) than those from primary responses such as early HIV infection (median = 5%, p = 0.0012) and primary hepatitis B vaccination (median = 2.8%, p = 0.0019) (**Fig. 2b**). Further, these influenza lineages had initial divergence levels similar to lineages from HIV patients first sampled >10 years after infection,^26,35^ and hepatitis B booster vaccination patients (**Fig. 2b**).^36^ Ebola virus infection, meningococcus vaccination, and early childhood development had median initial divergence levels of 3.5%, 6.6%, and 2.1%, respectively, but contained less than three adjusted measurably evolving lineages. Overall, these results are consistent with measurably evolving lineages from influenza vaccination arising mainly from pre-existing memory B cells.

While we found little measurable evolution in the blood following seasonal influenza vaccination, influenza vaccination has been shown to stimulate both naive and memory B cells to enter GCs.^3^ This raises the possibility that additional affinity maturation could be occurring in GCs, but its products are not enriched in the blood. Data from Turner, Zhou, Han et al. (2020)^3^ provided both blood samples and fine needle aspirations of lymph nodes (including GCs) from the same patient. By combining these samples, we identified 46 B cell lineages containing at least one GC B cell following influenza vaccination, and 107 lineages that contained none. To determine whether GC-engaged lineages were undergoing additional SHM, we tested whether they were enriched for measurable evolution. We found that 6.5% of lineages containing sequences from GC B cells were measurably evolving, compared to only 3.7% of lineages with no identified GC sequences. This signal of measurable evolution increased with the fraction of GC sequences. For instance, while 19% of lineages containing ≥10% GC sequences were measurably evolving, 50% of those with ≥25% GC sequences were (**Fig. 3a**). Lineages with higher proportions of GC sequences also had higher −log_10_(p values) from the date randomization test (linear regression slope p = 3.6×10^−7^, **Supplemental Fig. 7**). Measurably evolving lineages in this dataset did not contain significantly more sequences than other lineages, indicating these results were not confounded by lineage size (**Supplemental Fig. 8**). Finally, the measurably evolving lineages with the highest proportion of GC sequences contained monoclonal antibody sequences that bound to vaccine antigens (**Fig. 3b, c**). These lineages show signs of origin from memory B cells, such as clonal relatedness to blood plasmablasts sampled five days post-vaccination, and high mean germline divergence at their first sampled timepoint (6.3%, 7.2%, **Fig. 3b, c** respectively). Overall, these analyses demonstrate that influenza-binding B cell lineages are engaged by germinal centers and undergo additional, measurable evolution following vaccination.

**Fig. 3:**
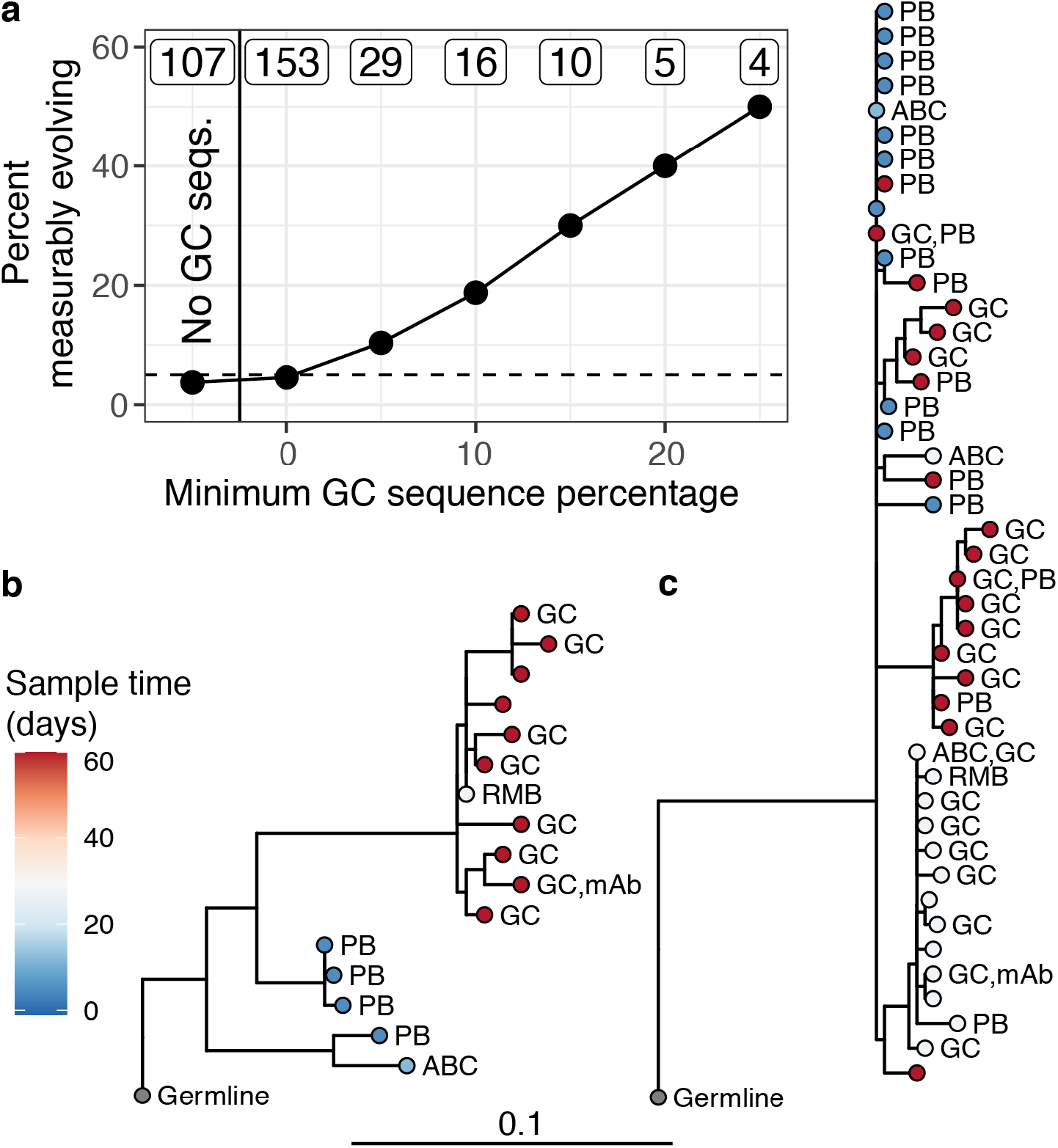
Germinal center engagement is positively related to measurable evolution following influenza vaccination. **a)** Percent of lineages that are measurably evolving given a minimum percentage of GC sequences. The minimum (inclusive) percent of GC sequences within a clone is shown on the x axis. The origin shows the percentage of measurably evolving lineages across all lineages. Left-most point shows lineages without any GC sequences. The total number of lineages in each category are listed above each point. The dashed line shows 5%, the expected false positive rate under the null hypothesis. **b-c)** Lineage trees showing measurably evolving lineages with the highest proportion of GC sequences. Tips are colored according to time collected post-vaccination and labeled by cell type if available. GC=germinal center, RMB = resting memory B, ABC = activated B cell, PB = plasmablast, and unlabeled tips are from bulk PBMC sequencing. mAb = influenza binding monoclonal antibody sequence (2018/2019 quadrivalent inactivated influenza virus vaccine).^3^ Branch lengths represent SHM/site, as shown by the shared scale bar.

Affinity maturation is a rapid evolutionary process. It is perhaps surprising that, while we identified conditions enriched for measurably evolving lineages, most lineages in circulation were not measurably evolving (**Table 1**). One explanation is that our analyses did not use sufficiently long sampling intervals to detect affinity maturation, though we believe this is unlikely. Experimental studies in mice have estimated that SHM occurs at ~10^−3^ SHM/site/division,^37^ and that GC B cells cycle every 6 – 12 hours.^7,38,39^ Simulations using conservative assumptions (strong selection, 24 hour cell cycle) and replicating the sample structure of our shortest-term influenza dataset (4 weeks), showed high power with >90% true positive rate (**Fig. 1h**). Further, we found an enrichment of blood-derived measurably evolving lineages after only 4 weeks in one study,^29^ and after 8 weeks corresponding to a known context of affinity maturation (GC entry, **Fig. 3**). Overall, these results show that the sample times of our surveyed datasets should be sufficient to detect ongoing SHM if it were occurring. A more plausible explanation for the lack of measurable evolution is that most lineages in the blood are either nonspecific to the condition being studied, or derive from a GC-independent response.^10,40,41^ Another explanation is that lineages may not remain in GCs continuously, which would slow the rate of evolution compared to our simulations.

The temporal dynamics of affinity maturation in humans following influenza vaccination are not well understood. This study demonstrates how a phylogenetic test of measurable evolution can be a powerful tool to detect ongoing SHM using longitudinally sampled BCR datasets across a wide array of immunological conditions, including influenza virus vaccine responses. While measurable evolution does not necessarily indicate affinity-driven selection, it does indicate ongoing SHM. It is hypothesized that seasonal influenza vaccination has poor efficacy because pre-existing memory B cells interfere with novel affinity maturation.^1^ Our results show that, indeed, there are few measurably evolving lineages in the blood following influenza vaccination. However, we also show that influenza vaccination is capable of stimulating measurable evolution in influenza-binding memory B cell lineages re-entering the GC. Thus, seasonal influenza virus vaccination in adults induces a GC reaction where maturation of vaccine-induced B cell lineages occurs, including those recruited from the pre-existing memory B cell compartment.

## Methods

### BCR sequence datasets and pre-processing

All longitudinally sampled B cell receptor (BCR) repertoire datasets were publicly available and obtained both from primary publications and through the Observed Antibody Space database^22^ (OAS; antibodymap.org, accessed 6/2020). Both assembled nucleotide sequences and deduplicated amino acid sequences were obtained from OAS. To reduce the effect of sequencing error in OAS datasets, only nucleotide sequences corresponding to an amino acid sequence with a redundancy of at least 2 were included. Datasets obtained from OAS are labeled in **Table 1**. Raw sequence data obtained from Nielsen et al. (2019)^28^ were pre-processed with pRESTO v0.5.13.^42^ Quality control was performed by first removing all sequences with a Phred quality score < 20, length < 300bp, or any missing (“N”) nucleotides. The 3’ and 5’ ends of each read were matched to forward and constant region primers with a maximum error rate of 0.1. The region adjacent to the constant region primer was exactly matched to sub-isotype specific internal constant region sequences. Only sequences with the same isotype predicted by their constant region primer and internal constant region sequence were retained. Identical reads within the same isotype were collapsed and sequences observed only once were discarded. All other datasets used processed BCR sequence data provided by the authors of their respective publications. Data from Wang et al. (2014)^43^ were processed in Hoehn et al. (2019).^44^ Data from Jiang et al. (2020b)^45^ used only blood samples.

### BCR sequence processing, genotyping and clonal clustering

Datasets were processed using the Immcantation framework (immcantation.org). V(D)J gene assignment on data obtained from OAS and Nielsen et al (2019)^28^ was performed using IgBLAST v1.13^46^ against the IMGT human germline reference database^47^ (IMGT/GENE-DB v3.1.24; retrieved August 3^rd^, 2019). V(D)J gene assignments and clonal cluster assignments were already available in all other datasets and were retained. Non-productively rearranged sequences were excluded. Using Change-O v1.0.0,^48^ the V and J genes of unmutated germline ancestors for each sequence were constructed with D segment and N/P regions masked by “N” nucleotides. Sequence chimeras were filtered by removing any sequence with more than 6 mutations in any 10 nucleotide window. Individual immunoglobulin genotypes were computationally inferred using TIgGER v1.0.0 and used to finalize V(D)J annotations.^49^ To infer clonal clusters, sequences were first partitioned based on common V and J gene annotations, and junction region length. Within these groups, sequences differing from one another by a specified Hamming distance threshold within the junction region were clustered into clones using single linkage hierarchical clustering.^17^ The Hamming distance threshold was determined by finding the local minimum of a bimodal distance to nearest sequence neighbor plot using SHazaM v1.0.2.999.^50^ In cases where automated threshold detection failed, usually because the distance to nearest neighbor distribution was not bimodal, the threshold was set to 0.1 and verified by manual inspection. Finally, the V and J genes of unmutated germline ancestors for each clone were constructed using masked D segments and N/P regions.

### Testing for measurable evolution

Testing for measurable evolution begins with building B cell lineage trees. Within each B cell clone, identical sequences or those differing only by ambiguous nucleotides were collapsed unless they were sampled at different timepoints. To reduce computational complexity, lineages were randomly down-sampled to at most 500 sequences each. B cell lineage tree topologies and branch lengths were estimated using maximum parsimony using the pratchet function of the R package *phangorn* v2.5.5.^51^ R packages *dowser* v0.0.3,^19^ *alakazam* v1.0.2.999,^48^ and *ape* v5.4-1^52^ were also used for phylogenetic analysis. Trees were visualized using *ggtree* v2.0.4^53^, and other figures were generated using *ggplot2* v3.3.2^54^ and *ggpubr* v0.4.0.^55^ R v3.6.1^56^ was used for all statistical analyses.

To test for measurable evolution over time, we use a modified version of the previously described phylogenetic date randomization test^14,15^ implemented in *dowser* v0.0.3.^19^ Briefly, for a given tree the divergence of each tip was calculated as the sum of branch lengths leading to the tree’s most recent common ancestor (MRCA). We next calculated the Pearson correlation between the divergence and sampling time of each tip. Measurably evolving lineages should show a positive correlation between divergence and time (**Fig. 1a**). We next identified monophyletic clades containing only sequences from a single timepoint (here referred to as “clusters”). We then randomly permuted sampling times among clusters, such that all sequences within each cluster had the same, randomly chosen timepoint. We next measured the correlation between divergence and time in this randomized tree, and repeated the process 100,000 times. We then estimated the p value that the observed correlation between divergence and time was no greater than expected from random distribution of times among clusters. This p value was calculated as the proportion of permutation replicates that had an equal or higher correlation than in the observed tree. We used a pseudocount of one for this calculation. The minimum possible p value for a lineage was calculated as one divided by possible number of distinct cluster permutations.

We modified the data randomization test to account for the high degree of topological uncertainty of many B cell lineage trees. More specifically, B cell lineage trees often contain large clusters of zero-length branches (soft polytomies) that represent high uncertainty in branching order (e.g. **Supplemental Fig. 1**). In bulk BCR data these polytomies are likely due to PCR error or sequencing error. If polytomies are resolved randomly into bifurcations, this can produce more single timepoint monophyletic clades than necessary (**Supplemental Fig. 1**) and lead to a high false positive rate of the date randomization test (**Supplemental Fig. 2**). To ensure this source of uncertainty did not increase the false positive rate of our analyses, we resolved bifurcations within each polytomy such that sequences from the same timepoint were grouped in the fewest possible number of single-timepoint monophyletic clades before performing permutations.

The clustered date randomization approach is more conservative than tests that permute tips uniformly,^57^ but has been shown to be less biased if different sub-populations are sampled at each timepoint.^15^ To explore the effect of this modelling choice, we repeated the analyses in **Table 1** using two-tailed clustered and uniform date randomization tests (**Supplemental Fig. 2**). Two-tailed tests can identify lineages with a significant positive or negative correlation between divergence and time. This is useful because a significant negative correlation between divergence and time is biologically implausible and represents a likely false positive result. Due to multiple testing under an alpha value of 0.025, we expect no more than 2.5% of lineages to have a significant negative correlation from these two-tailed tests. We found the uniform permutation test had a high rate of negatively evolving lineages (mean = 8.3%), indicating a high false positive rate. By contrast, the clustered permutation test without resolved polytomies had a mean rate of only 2.2% negatively evolving lineages, approximately as expected given an alpha value of 0.025. Resolving polytomies and then performing the clustered permutation test improved performance even more, with a mean rate of 1.2% negatively evolving lineages and no dataset having more than 2.8% of lineages negatively evolving. This analysis shows the uniform date randomization test is prone to false positives in empirical B cell data, while the clustered date randomization test with resolved polytomies corrects this issue. All other tests performed in this study used a one-tailed, clustered date randomization test with resolved polytomies and an alpha value of 0.05.

To identify and characterize measurably evolving lineages while adjusting for multiple testing, all lineages tested were pooled together and p values were adjusted using the Benjamini-Hochberg procedure^34^ implemented in the R v3.6.1 function p.adjust.^56^ Lineages with adjusted p values < 0.1 were referred to as adjusted measurably evolving lineages (**Fig. 2b**).

It is possible that the results reported are affected by the size (number of sequences) of lineages in each dataset. A large number of lineages without adequate power could result in a spurious lack of measurable evolution. To ensure the lineages included in each study were adequately powered, we included only lineages with at least 15 sequences, were sampled over at least three weeks, and had a minimum possible p value < 0.05 based on the number of distinct permutations of timepoints among clusters. If measurable evolution were still strongly confounded by lineage size even after these filtering steps, we would expect measurably evolving lineages to be larger on average than non-measurably evolving lineages. By contrast, measurably evolving lineages were significantly larger than non-measurably evolving lineages in only 5/21 datasets surveyed (**Supplemental Fig. 8**), indicating our results are not strongly confounded by lineage size.

### Simulation-based power analysis

We used simulations to determine whether the clustered date randomization test was sufficiently powered to detect ongoing affinity maturation. These analyses used the *bcr-phylo* package accessed 9/21/2020,^20,21^ which simulates clonal lineages of B cells undergoing affinity maturation against a target sequence. For all simulations, a random naive heavy chain sequence was chosen from those provided in *bcr-phylo* and the rate of SHM was set to the default of λ=0.356, which corresponds to an SHM rate of ~ 0.001 SHM/site/division.^9^ Mutations were introduced according to the S5F model.^50^ Selection strength was chosen to be either 0 (neutral), or 1 (entirely affinity driven). A single target sequence was chosen for affinity maturation. All other parameters were set to their default.

We performed two sets of simulations. In the first, we simulated single B cell lineages from which 50 cells were sampled at generation 10, and 50 more cells were sampled after a specified number of additional generations (**Fig. 1g, Supplemental Fig. 3**). In the second type of simulation, we replicated the sampling strategy of Laserson et al. (2014)^13^. Briefly, for each clone in subject *hu420143* from Laserson et al. (2014)^13^, we simulated one lineage with the same number of cells (if enough cells had been generated) sampled after the number of generations corresponding to 1, 3, 7, 14, 21, and 28 days (**Supplemental Fig. 4**). The number of generations corresponding to each sample day was calculated using a strict generation time of either 12 hours or 24 hours, which are conservative given previous GC cycle estimates of 6 – 12 hours.^7,38,39^ These simulations used a selection strength of 1, which gave more conservative results in previous simulations (**Supplemental Fig. 3**).

To account for possible issues with clonal clustering, we did not preserve clonal identities among simulated sequences in either simulation type. Instead, we pooled sequences from all simulation repetitions under a particular parameter set and used the same clonal clustering method used for empirical data analyses to group them into clonal clusters. We did not repeat the genotyping or chimera filtering steps done on empirical data analyses as genotyped individuals and sequence chimeras were not part of the simulations. We performed the clustered date randomization test with resolved polytomies on each lineage with a minimum possible p value < 0.05. Because all sequences were simulated under affinity maturation, the proportion of lineages with p < 0.05 indicated the true positive rate of the test. To determine the false positive rate, we randomized sample times among tips within each tree and repeated the date randomization test (**Supplemental Figs. 3-4**). Here, the proportion with p < 0.05 indicated the false positive rate.

Scripts to reproduce all analyses performed are available at: https://bitbucket.org/kleinstein/projects/src/master/Hoehn2021_biorxiv

## Acknowledgements

We would like to thank Dr. Louis Du Plessis for helpful discussion, and Dr. Julian Q. Zhou for providing processed data. This work was funded in part by National Institutes of Health, National Institute of Allergy and Infectious Diseases grant R01 AI104739, and by the European Research Council under the European Union’s Seventh Framework Programme (FP7/2007-2013)/European Research Council grant agreement number 614725-PATHPHYLODYN. The Ellebedy laboratory was supported by NIAID grants R21 AI139813, U01 AI141990, and NIAID Centers of Excellence for Influenza Research and Surveillance (CEIRS) contract HHSN272201400006C to A.H.E. J.S.T. was supported by NIAID 5T32CA009547.

## Author contributions

K.B.H. and S.H.K designed the study and composed the manuscript. K.B.H performed analyses. K.B.H and F.I.M. developed data processing pipelines. A.H.E, J.S.T., and R.J. helped interpret the data and provided processed datasets. O.G.P. advised on phylogenetic methods. All authors reviewed the manuscript.

## Competing interests

K.B.H. receives consulting fees from Prellis Biologics. S.H.K. receives consulting fees from Northrop Grumman. The Ellebedy laboratory received funding under sponsored research agreements from Emergent BioSolutions and AbbVie.

