## Supplemental Figures 1 - 8 for "Human B cell lineages engaged by germinal centers following influenza vaccination are measurably evolving"

### Human B cell lineages engaged by germinal centers following influenza vaccination are measurably evolving

Kenneth B. Hoehn, Jackson S. Turner, Frederick I. Miller, Ruoyi Jiang, Oliver G. Pybus, Ali H. Ellebedy, Steven H. Kleinstein

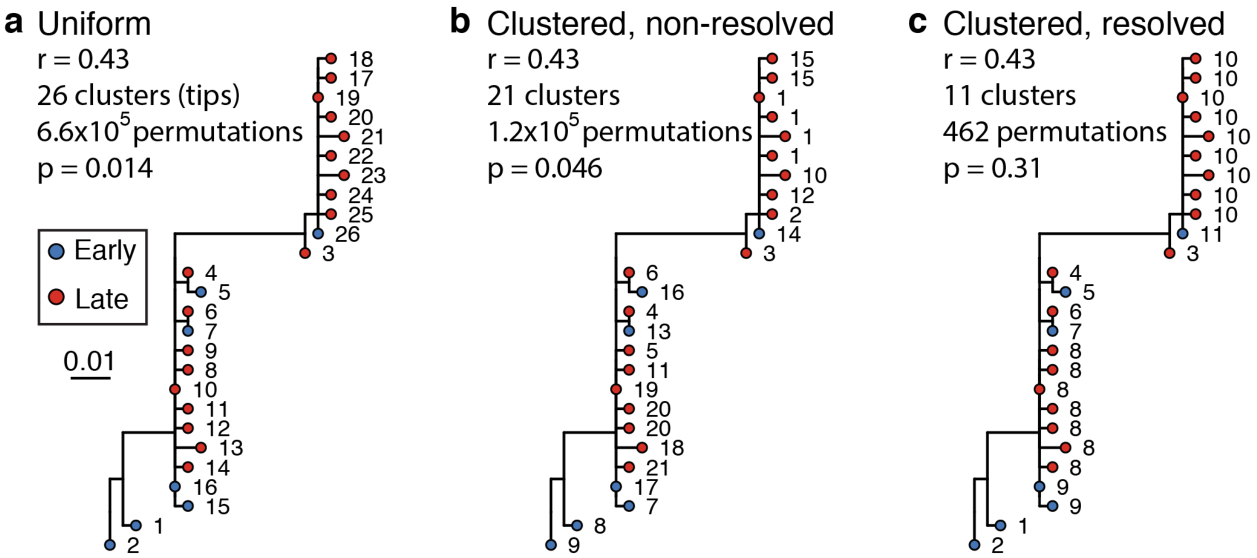


**Supplemental Fig. 1:** Comparison of date randomization tests on an example tree from Galson et al. (2015b)[1] showing little evidence of ongoing SHM. This tree contains two large polytomies consisting of multiple short branches radiating out from a central node. These features can result from sequencing error or PCR error in bulk BCR data, where errors create spurious, unique sequences one mutation away from a single real sequence. Permuting these tips uniformly among each other leads to these spurious tips being treated as independent data points, and can lead to high false positive rates if not corrected (**Supplemental Fig. 2**). While visual inspection of this tree shows little evidence of increase in SHM over time, it has a date randomization test $p < 0.05$ unless its polytomies are resolved.

In the panels above, each tip is a sequence labeled with its cluster assignment. **a)** Tips are permuted individually, meaning each tip is a separate cluster. This leads to $6.6 \times 10^5$ distinct permutations of time labels along the tree, and a $p < 0.05$. **b)** Tips belonging to single-timepoint monophyletic clades are grouped into clusters, equivalent to Murray et al. (2016).[2] Timepoints are permuted among these clusters, which reduces the number of possible permutations. This also reduces the significance of the relationship between divergence and time. However, because the polytomies are randomly resolved into bifurcations with zero-length branches, each polytomy has multiple clusters with the same timepoint. For instance, clusters 1, 15, 10, 12, and 2 could be grouped in the same cluster but are kept distinct. **c)** Bifurcations using zero-length branches within the polytomies are rearranged to give the fewest possible number of monophyletic single-timepoint clusters. Resolving polytomies effectively treats same-timepoint sequences within polytomies as the same data point, appropriately showing this tree does not have sufficient evidence of measurable evolution.

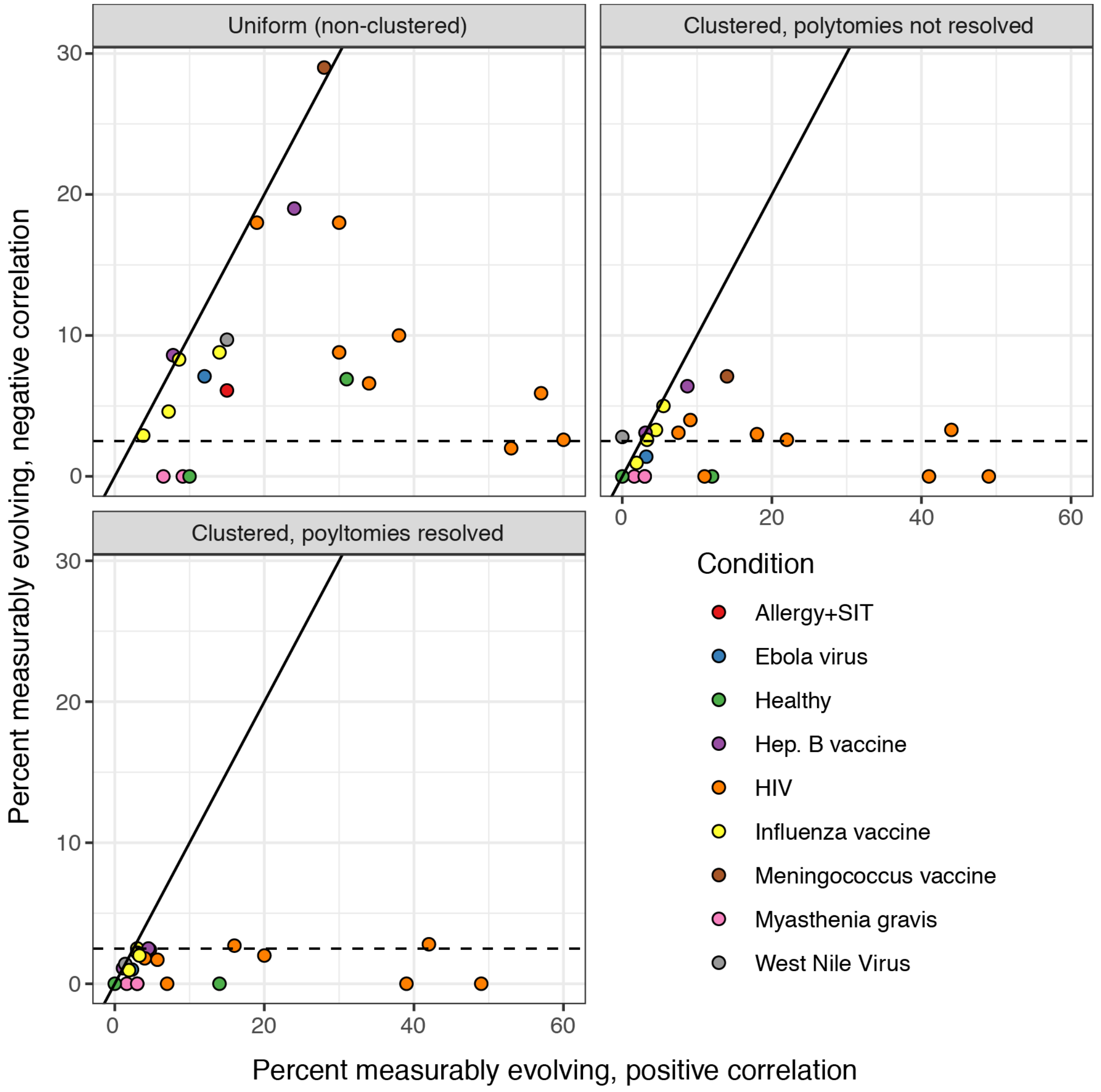


**Supplemental Fig. 2**: Uniform vs. clustered date randomization test performance. The date randomization test can be performed using either uniform permutations, in which the timepoint of each tip is permuted separately, or clustered permutations, in which timepoints are permuted among single-timepoint monophyletic clusters. It can also be performed using clusters after polytomies have been resolved into the smallest possible number of single-timepoint clades. Using a two-tailed test, we determine whether a lineage is positively measurably evolving (correlation between divergence and time > 0, $p < 0.025$) or negatively measurably evolving (correlation < 0, $p < 0.025$). Measurable negative evolution indicates decreasing divergence over time, which is biologically implausible and likely represents false positives. This could be due to population structure at different timepoints. See Murray et al (2016).[2]

In the panels above, we repeated the analyses in **Table 1** using two-tailed tests with each permutation strategy. The x axis shows the percent of positively measurably evolving lineages for each study, while the y axis shows the percent of negatively measurably evolving lineages, which are interpreted as false positives. The dashed line shows 2.5%, the maximum expected percent of negatively evolving lineages. Only clustered permutations with resolved polytomies – used in all other analyses in this manuscript – fully controlled this error metric.

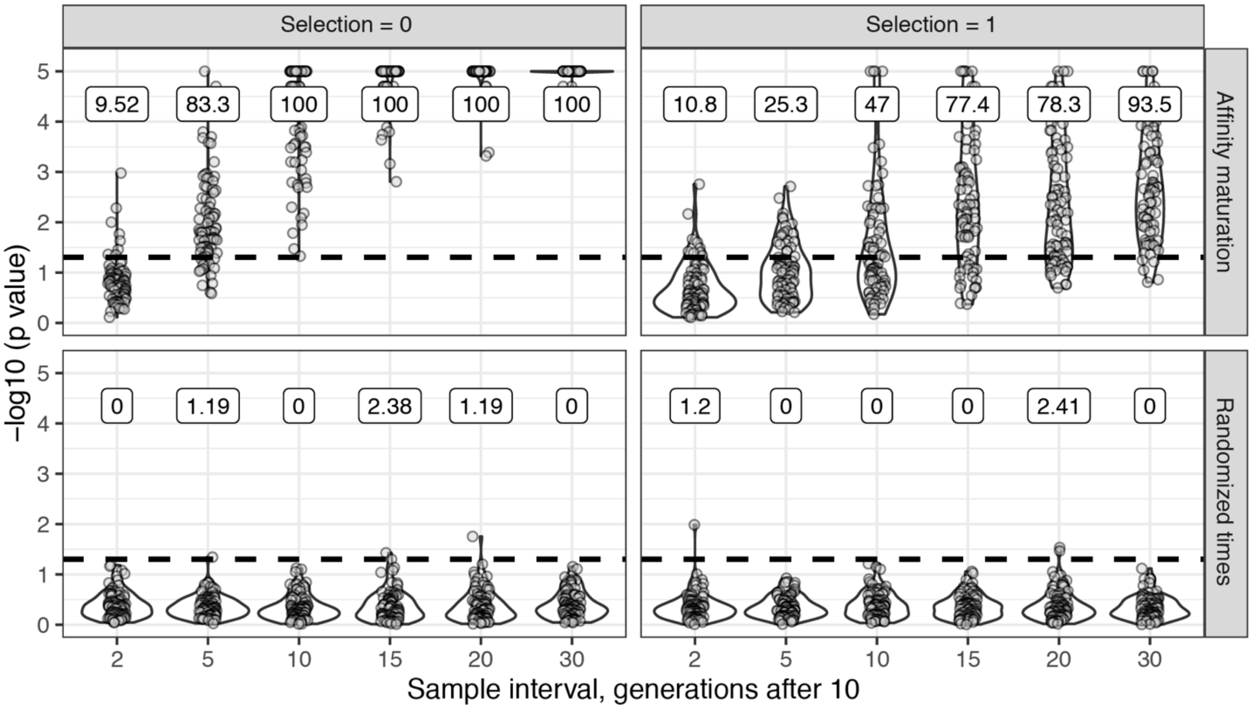


**Supplemental Fig. 3**: Affinity maturation simulations. For upper panels (Affinity maturation) each lineage was simulated for 10 GC cycles before 50 cells were sampled, if available. Affinity maturation continued for the specified number of additional GC cycles (x axis) before a second sampling of 50 cells. This process was repeated for 100 repetitions for the specified number of GC cycles, and given the specified strength of selection. Selection = 0 corresponds to neutral evolution, while Selection = 1 corresponds to strong selection for matching to a single target sequence. Default parameters from *bcr-phylo*[3,4] were used otherwise. The y axis shows the -log10(p value) for the date randomization test, with dots above the horizontal dashed line representing measurably evolving lineages ($p < 0.05$). The percentage of measurably evolving lineages for each set of simulations is shown above the dashed line, rounded to three significant digits. Only simulated lineages with a minimum possible $p < 0.05$ were tested in simulations. Because all lineages were undergoing affinity maturation in the upper panels, this corresponds to the true positive rate. Lower panels (Randomized times) show results from the same simulated data but with sampling times randomized among sequences. Because these simulations are effectively not evolving over time, the numbers above show the false positive rate in the lower panels.

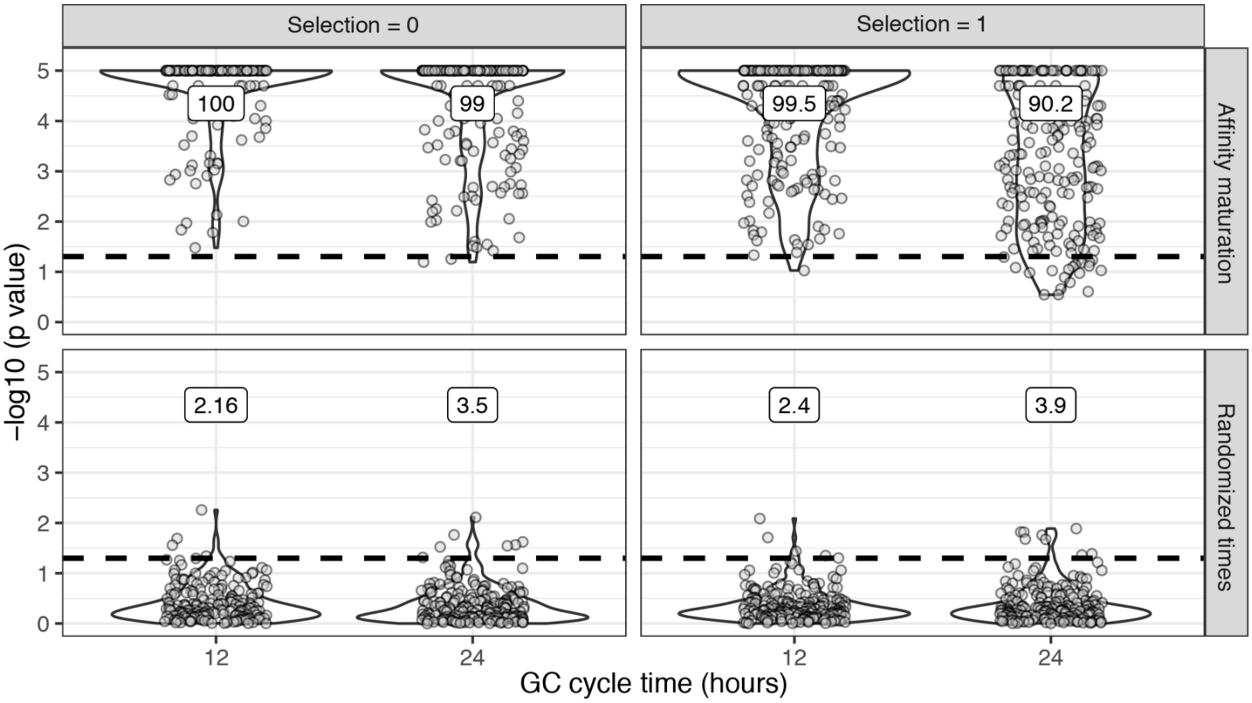


**Supplemental Fig. 4**: Simulations based on an empirical dataset. Simulations were performed to replicate the sampling strategy of Laserson et al (2014)[5,6], in which an individual was sampled at 6 timepoints between 1 day and 28 days following influenza vaccination. We excluded pre-vaccination samples as well as the sample taken 1 hour after vaccination because it was too early for any GC cycles to occur in simulations. For each simulation, we selected a lineage *C* from subject *hu420143*. To calculate the number of GC cycles to simulate, we divided the sample times (hours post vaccination) of lineage *C* by the specified GC cycle time (x axis). We then simulated affinity maturation as in **Supplemental Fig. 3**, and sampled the same number of cells as were present in *C* at the corresponding time. We repeated this process for each lineage in subject *hu420143* with at least 15 sequences sampled over three weeks and a minimum possible p value $< 0.05$. The percentage of measurably evolving lineages for each set of simulations is shown above the dashed line, rounded to three significant digits. Only simulated lineages with a minimum possible $p < 0.05$ were tested in simulations. Because all lineages in the upper panels (Affinity Maturation) were undergoing affinity maturation, this corresponds to the true positive rate. Lower panels (Randomized times) show results from the same simulated data but with sampling times randomized among sequences. Because these simulations are effectively not evolving over time, the numbers above show the false positive rate in the lower panels.

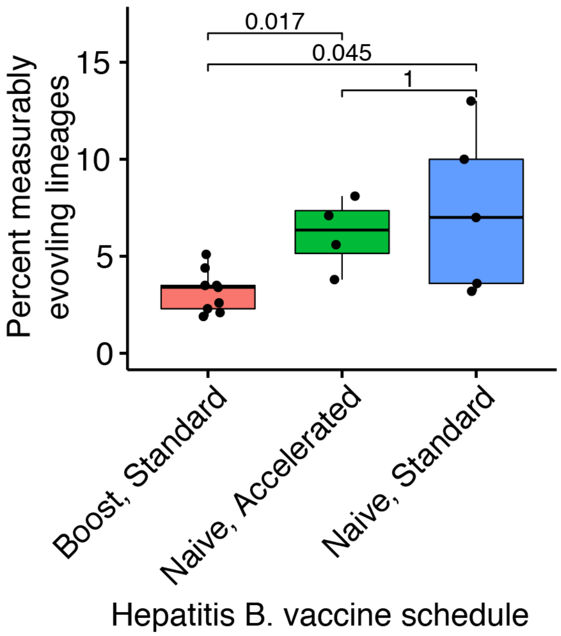


**Supplemental Fig. 5**: Measurably evolving lineages under different hepatitis B vaccine contexts. Hepatitis B booster vaccine data was obtained from Galson et al (2015a)[7] and consisted of nine previously vaccinated patients sampled 4 times between 0 and 28 days after a single vaccination. Hepatitis naive data were obtained from Galson et al. (2016).[8] These patients were all vaccine-naive, were given 3 vaccinations, and sampled at 7 timepoints. Five patients received "standard" vaccinations at days 0, 28, and 168, and were sampled at days 0, 7, 28, 35, 168, 175, and 208. Four patients received "accelerated" vaccinations at days 0, 28, and 56, and were sampled at days 0, 7, 28, 35, 56, 63, and 96. P values were calculated using a Wilcoxon test.

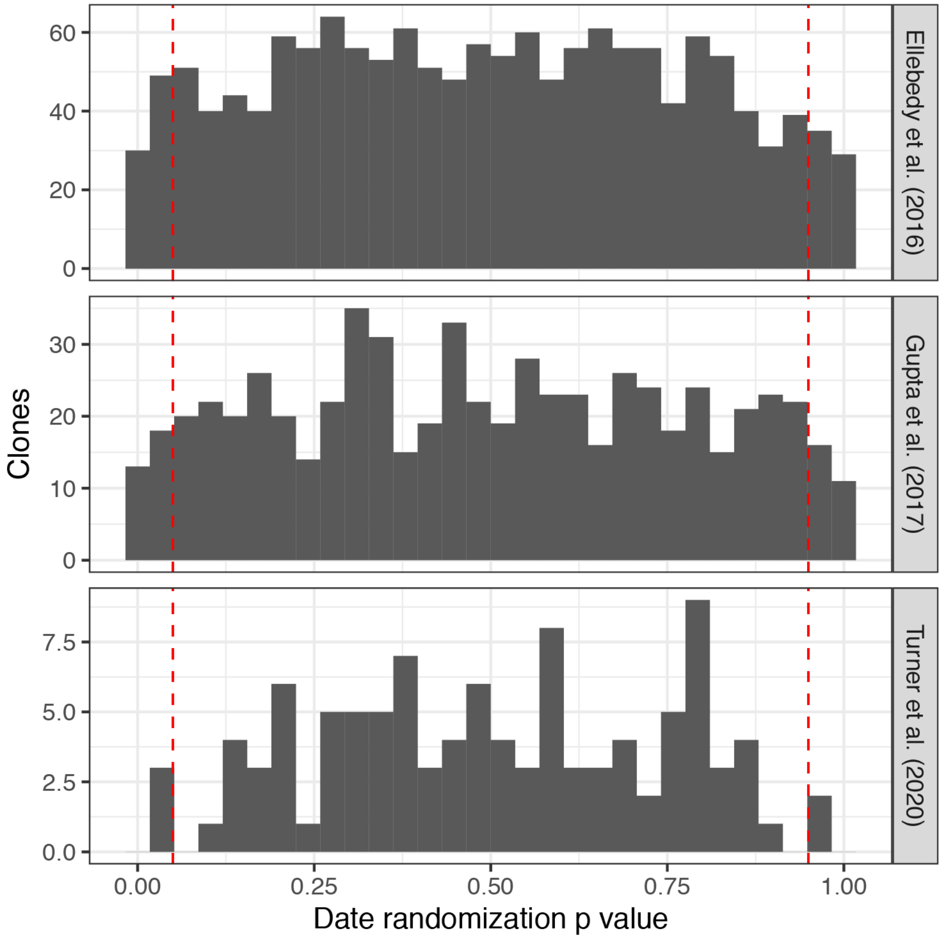


**Supplemental Fig. 6**: Date randomization test p value histograms from blood-derived lineages in three influenza studies.

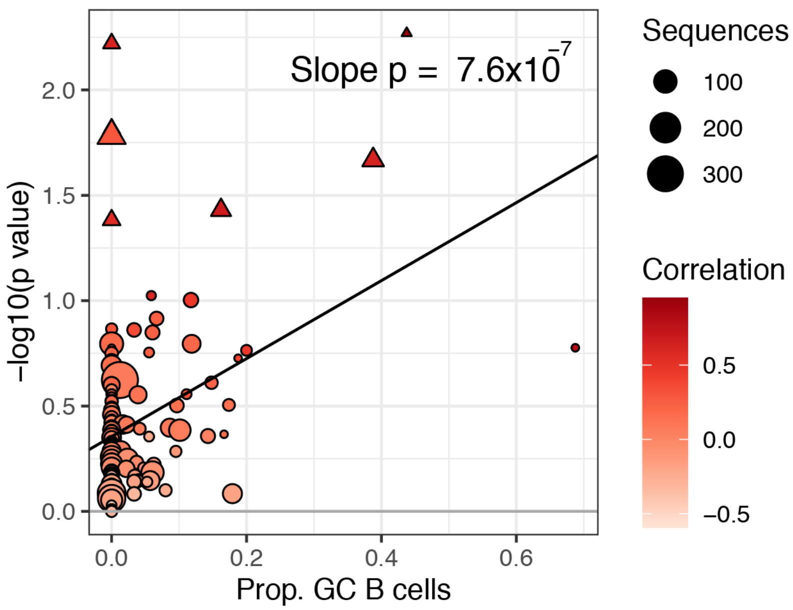


**Supplemental Fig. 7:** Germinal center engagement is positively related to measurable evolution following influenza vaccination. Proportion of GC sequences within a lineage is positively related to low date randomization test p value; i.e., high -log10(p value). Lineages with $p < 0.05$ are shown as triangles. Points are colored by correlation between divergence and time.

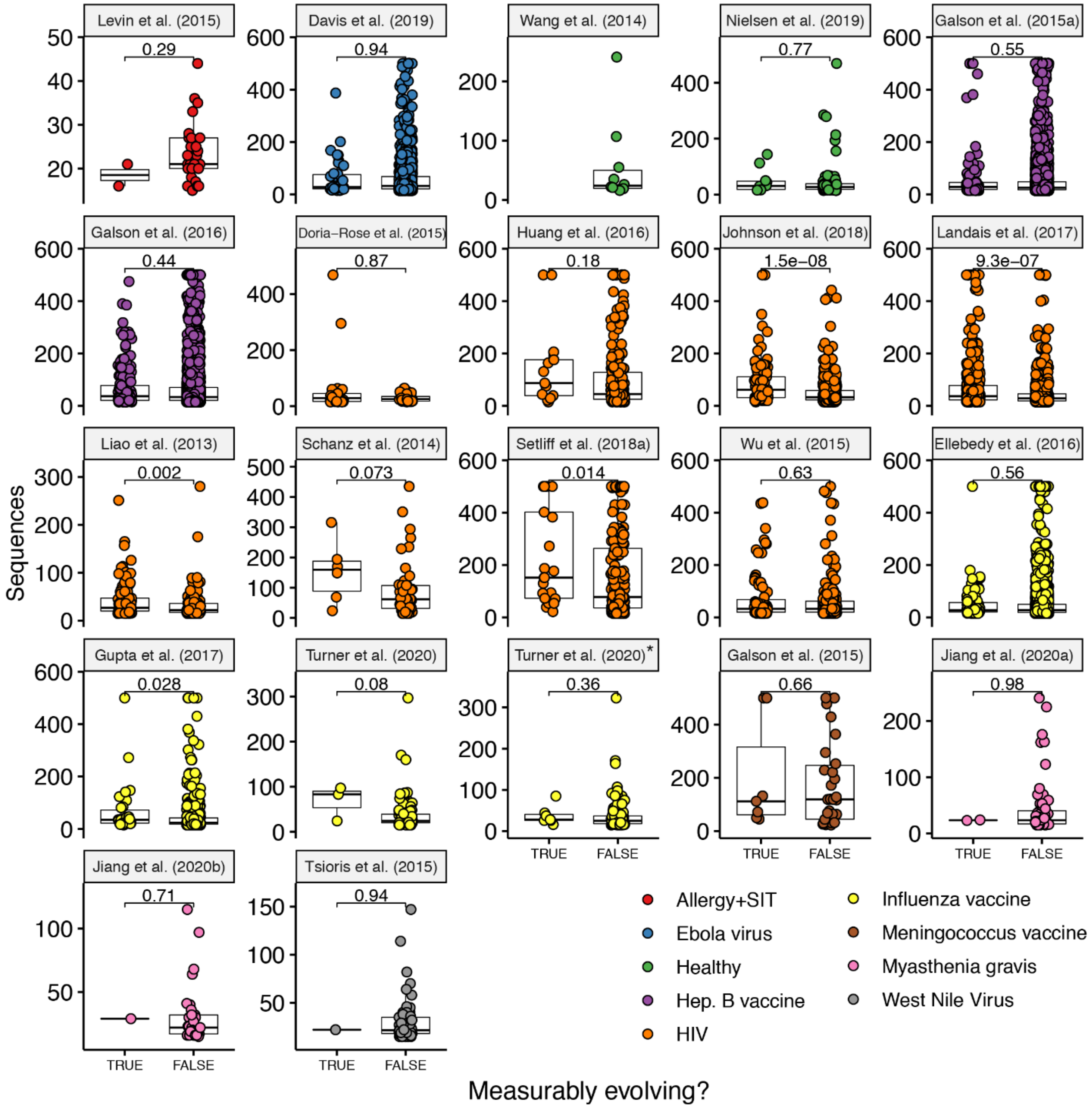


**Supplemental Fig. 8:** Number of sequences per lineage in measurably evolving vs non-measurably evolving lineages. See **Table 1** for details on each study. P values are computed using a Wilcoxon test. Turner et al. (2020)* included all samples (blood and fine needle aspiration) while Turner et al. (2020) included only blood samples.
